# A *Csf1r* lineage gives rise to dermal lymphatic endothelial cells

**DOI:** 10.64898/2026.03.17.712362

**Authors:** Giovanni Canu, Rosamaria Correra, Alice Plein, Laura Denti, Alessandro Fantin, Christiana Ruhrberg

## Abstract

Lymphatic vessels are formed during embryonic and postnatal development to facilitate interstitial fluid clearance and immune regulation after birth. Their organ-specific heterogeneity in organisation and function is preceded by heterogenous origins of lymphatic endothelial cells (LECs), the main building blocks of lymphatic vessels. In the dermis, a subset of LECs was reported to arise from blood capillaries, which themselves differentiate, in part, from paraxial mesoderm. However, it is not known whether additional cell lineages contribute to the dermal LEC population. Here, we have combined transcriptomic analyses with genetic lineage tracing and wholemount immunostaining to show that 60% of LECs in the embryonic day (E) 13.5 and E15.5 dermis are derived from a cell lineage that expresses *Csf1r*, a marker of myeloid cells and their progeny. *Csf1r* lineage LECs persist in adult dermal lymphatic vasculature and are indispensable for normal lymphatic development, because *Prox1* deletion within the *Csf1r* lineage causes dermal oedema and blood-filled lymphatic vessels. As *Csf1r* lineage dermal LECs do not themselves express *Csf1r* and also do not arise from *Csf1r-*expressing differentiated myeloid cells, our findings imply the existence of a *Csf1r*-expressing non-LEC precursor population for the majority of dermal LECs and will prompt further work to identify this cell population.

## INTRODUCTION

The lymphatic vasculature is essential for maintaining tissue-fluid balance, absorbing lipids and supporting immune surveillance. Accordingly, lymphatic dysfunction causes lymphoedema, tissue fibrosis, subcutaneous fat accumulation and susceptibility to infections (Alitalo, 2011; Martin-Almedina, Mortimer and Ostergaard, 2021). To understand the formation of lymphatic vasculature during embryonic and postnatal development, much effort is currently directed at defining the cell lineages that gives rise to lymphatic endothelial cells (LECs), the main cellular constituent of lymphatic vasculature (Jafree *et al*., 2021).

Initial lineage tracing studies from the promoter of the *Tie2* gene demonstrated that LECs arise during embryogenesis from a population of blood vascular endothelial cells (BECs) in the cardinal vein from embryonic day (E) 9.5 onwards in the mouse (Srinivasan *et al*., 2007). Following their differentiation from BECs, venous-derived LECs protrude and delaminate from the cardinal vein to assemble into the primordial jugular lymph sacs (François *et al*., 2012; Yang *et al*., 2012). Subsequent work revealed that organ LECs have multiple, organ-specific origins (Jafree *et al*., 2021; Oliver, 2022). For example, lineage tracing from the *Isl1* promoter was used to demonstrate a second heart field origin of half of LECs in the ventral heart (Maruyama *et al*., 2019; Lioux *et al*., 2020). By contrast, lineage tracing with a temporally inducible *Kit* promoter suggested that a subset of LECs in the mesentery arises from haemogenic BECs (Stanczuk *et al*., 2015). A haemogenic origin was also proposed for a subset of LECs in the heart based on lineage tracing with the *Pdgfrb*-, *Vav1*- and *Csf1r* promoters (Klotz *et al*., 2015). By contrast, dermal LECs were reported to arise from E13.5 onwards in the mouse from BECs in the blood vascular capillary plexus (Pichol-Thievend *et al*., 2018). Complementary lineage tracing from the *Pax3* gene, which is a characteristic marker of paraxial mesoderm, identified BECs arising from paraxial mesoderm (Mayeuf-Louchart *et al*., 2014) and also a LEC subset in the dermis as well as in the heart (Stone and Stainier, 2019). However, it has not yet been investigated whether the *Csf1r* lineage, which also gives rise to BECs (Plein *et al*., 2018), additionally contributes LECs to the dermis.

Here, we have combined transcriptomic analyses with genetic lineage tracing and wholemount immunostaining to show that a cell lineage expressing the haematopoietic marker *Csf1r* gives rise to a large proportion of LECs in the dermis, and that their genetic ablation causes dermal oedema and erythrocyte entry into the lymphatic vasculature.

## MATERIALS AND METHODS

### Transcriptomic analysis

Raw bulk RNAseq reads of endothelial cells isolated from E12.5 *Csf1r-iCre*;*Rosa*^*tdTom*^ whole embryo (NCBI Gene Expression Omnibus database, accession number GSE117978) (Plein *et al*., 2018) were pre-processed to trim poor quality base calls and adaptor contamination using Trimmomatic v.0.36.4 (Bolger, Lohse and Usadel, 2014) and aligned to the mouse mm10 genome with STAR v.2.5.2b (Dobin *et al*., 2013). Mapped reads were deduplicated to reduce PCR bias using Picard v2.7.1.1 software (http://broadinstitute.github.io/picard/), and the reads-per-transcript were then calculated using FeatureCount v1.4.6.p5 software (Liao, Smyth and Shi, 2014). Differential expression was performed using the BioConductor package DESeq2 via the SARTools wrapper v1.3.2.0 (Varet *et al*., 2016).

### Animal procedures and mouse strains

Animal procedures were performed with Animal Welfare Ethical Review Body (AWERB) and UK Home Office approval. The following mouse strains were interbred: *Csf1r-iCre* (Deng *et al*., 2010), *Rosa*^*tdTom*^ (Madisen *et al*., 2010), *Spi1*^*+/-*^ (McKercher *et al*., 1996), *Rosa*^*Yfp*^ (Srinivas *et al*., 2001), *Csf1r-Egfp* (Burnett *et al*., 2004), *Csf1r-Mer-iCre-Mer* (Qian *et al*., 2011) and *Prox1*^*fl(Egfp)/+*^ (Iwano *et al*., 2012). All mice were maintained on a genetic background that was mostly C57BL/6J, with some contribution from 129/sv. To obtain mouse embryos of defined gestational age, mice were paired in the evening, and the presence of a vaginal plug the following morning defined as E0.5. For tamoxifen induction of CRE activity in mice carrying the *Csf1r-Mer-iCre-Mer* transgene, 1 mg tamoxifen (Sigma, dissolved in corn oil) was administered via a single intraperitoneal injection into each pregnant dam. Embryos were collected at the indicated times after culling of the pregnant dam by cervical dislocation. Embryos were fixed in 4% formaldehyde and washed in phosphate-buffered saline (PBS).

### Tissue immunostaining

Embryonic dorsal dermis was dissected from formaldehyde-fixed embryos and then incubated in PBS containing 2% serum-free protein block (DAKO), 2% bovine serum albumin and 0.4% Triton X-100 before staining with a combination of the following primary antibodies: goat anti-mouse NRP2 (R&D Systems #AF567, 1:100), rabbit anti-mouse PROX1 (Biolegend # 925202, 1:50), goat anti-mouse PROX1 (R&D Systems # AF2727, 1:100), rabbit anti-mouse LYVE1 (Angiobio # 11-034, 1:100), rat anti-mouse PECAM1 (BD Pharmigen # 553370, 1:50), rat anti-mouse EMCN (Santacruz # sc-65495, 1:50), goat anti-mouse FLT4 (R&D Systems # AF743, 1:100), rat anti-mouse TER119 (Biolegend # 116241, 1:100), rabbit anti-mouse GFP (MBL # 598, 1:200), goat anti-mouse TOM (Origene #AB1140-100, 1:250), or rat anti-mouse TOM (Chromotek #5F8, 1:200). After washing in PBS, we used appropriate Fab fragments as secondary antibodies: Alexa Fluor 647 donkey-anti goat (Stratech # 705-607-003, 1:200), Cy3 donkey anti-goat (Stratech #705-166-147, 1:200), Alexa Fluor 488 donkey anti-rabbit (Stratech #711-547-003, 1:200), Alexa Fluor 647 donkey-anti rabbit (Stratech # 711-607-003, 1:200), Alexa Fluor 488 donkey anti-rat (Stratech # 712-547-003, 1:200), or Cy3 donkey anti-rat (Stratech # 712-166-150, 1:200). DAPI-counterstained samples were imaged on a LSM710 confocal microscope (Zeiss).

## RESULTS

### Transcriptomic analysis and immunostaining identify *Csf1r* lineage traced LECs

Prior analysis of a bulk RNAseq transcriptomic dataset (GSE117978) from E12.5 *Csf1r-iCre*;*Rosa*^*tdTom*^ whole embryos had shown that liver sinusoidal BEC markers such as *Mrc1* and *Lyve1* were over-represented in *Csf1r* lineage ECs (TOM+) compared to other ECs (TOM-) (Plein *et al*., 2018). As *Lyve1* was originally identified also as a lymphatic marker (Banerji *et al*., 1999), we re-examined this dataset also for the expression of other known LEC markers, including *Prox1*, a transcription factor which is required for LEC specification, *Flt4*, which encodes the VEGFR3 receptor for VEGFC, a growth factor that promotes LEC proliferation and migration, and *Nrp2*, which encodes a VEGFR3 co-receptor (Veikkola *et al*., 2001; Wigle *et al*., 2002; Yuan *et al*., 2002). Similar to *Lyve1*, the transcripts from all three LEC marker genes were enriched in *Csf1r* lineage traced (TOM+) ECs compared to lineage negative (TOM-) ECs (**Fig. 1A,B**). The *tdTomato* transcript was enriched in TOM+ ECs, as expected, whereas the *Csf1r* transcript was not detected in either EC population (**Fig. 1C**), as previously reported (Plein *et al*., 2018), indicating that these ECs do not express *Csf1r* themselves but arise from a *Csf1r+* progenitor. Quantitative reverse transcriptase (qRT)-PCR analysis of TOM+ versus TOM-ECs isolated from a different cohort of E12.5 *Csf1r-iCre*;*Rosa*^*tdTom*^ whole embryos confirmed that LEC markers were significantly enriched in the TOM+ EC population (**Fig. 1D**).

**Fig 1.**
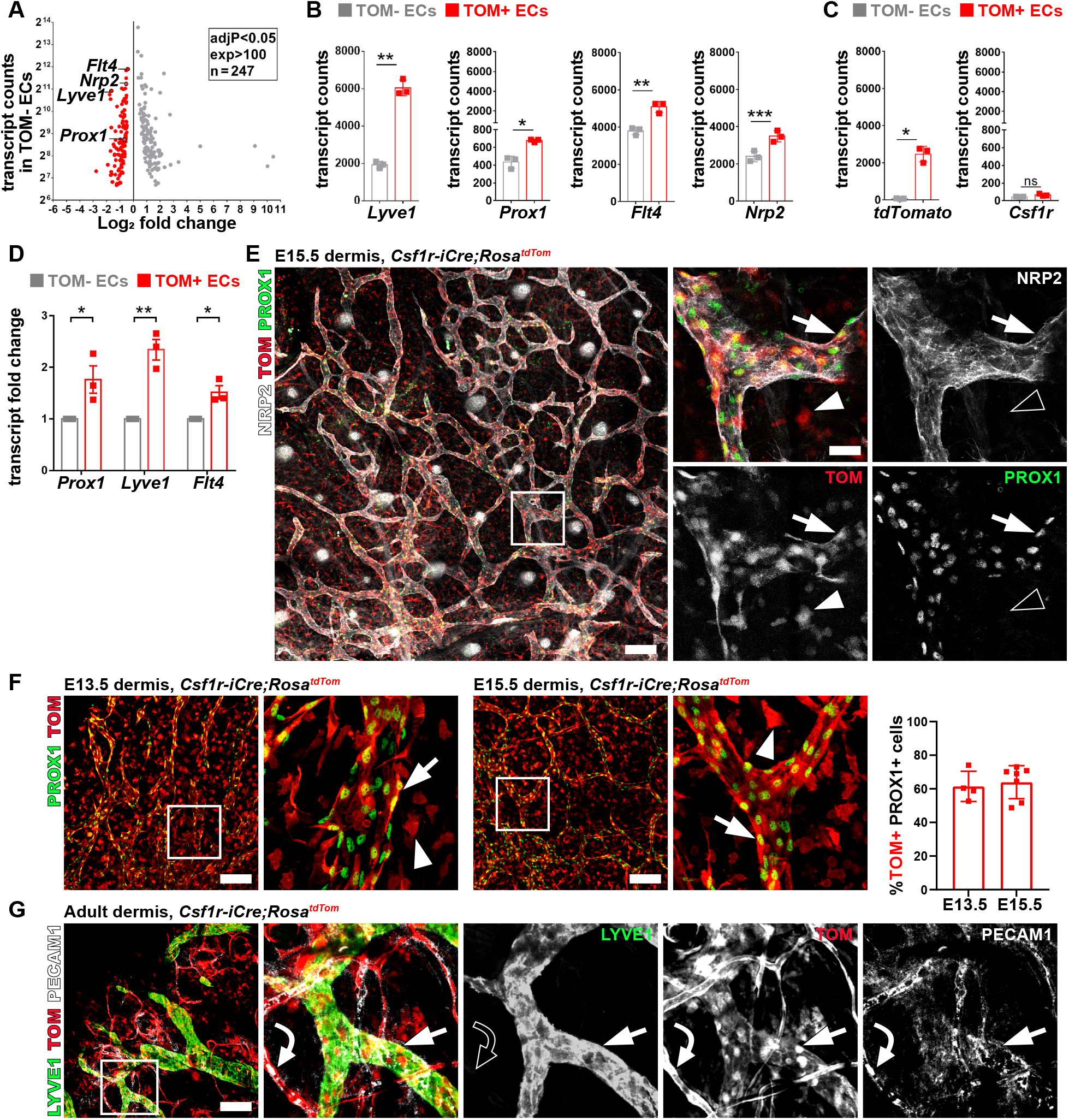
*Csf1r* lineage contribute lymphatic endothelial cells in the mouse dermis. **A-C** Bulk RNA-seq analysis of tdTomato (TOM)+ and TOM-endothelial cells (ECs) from whole E12.5 *Csf1r-iCre;Rosa*^*tdTom*^ mouse embryos (dataset GSE117978). **A** Volcano plot of significantly differentially expressed transcripts with more than 100 counts per transcript; relevant genes are named; grey and red data points represent transcripts with at least twofold over- or under-representation, respectively (*i*.*e*., red data points are over-represented and grey data points under-represented in TOM+ ECs compared to TOM-ECs). **B**,**C** Relative expression levels for markers typical of lymphatic endothelial cells (LECs) (**B**) or *tdTomato* and *Csf1r* (**C**); mean ± SD, n = 3 embryos; NS, not significant, *P < 0.05, **P < 0.01 (Benjamini–Hochberg’s multiple comparisons test for P value adjustment, Padj). **D** RT-qPCR expression analysis for the indicated genes relative to *Actb*, showing fold change expression in TOM+ ECs relative to TOM-ECs; mean ± SD, n = 3 embryos; *P < 0.05, **P < 0.01 (two-tailed unpaired t-test). **E** Immunofluorescence staining with the indicated markers of E15.5 *Csf1r-iCre*;*Rosa*^*tdTom*^ dorsal dermis (scale bar: 200 μm); the square indicates an area shown at higher magnification in the adjacent panels (scale bar: 25 μm), and shown for the different markers also in grey scale. **F** Immunofluorescence staining with the indicated markers for quantification of TOM+ LECs in E13.5 and E15.5 *Csf1r-iCre*;*Rosa*^*tdTom*^ dermis (scale bars: 100 μm); the squares indicate areas shown at higher magnification in the adjacent panels. The bar plot shows the fraction of TOM+ PROX1+ cells in E13.5 and E15.5 *Csf1r-iCre*;*Rosa*^*tdTom*^ dermis. Mean ± SD, n = 4 embryos; each dot represents the value for one embryo. **G** Immunofluorescence staining with the indicated markers of adult *Csf1r-iCre*;*Rosa*^*tdTom*^ ear dermis (scale bar: 100 μm). The square indicates an area shown at higher magnification in the adjacent panel, and shown for the different markers also in grey scale. Arrows indicate TOM+ LECs; arrowheads indicate TOM+ macrophages; curved arrows indicate TOM+ BECs; empty symbols indicate lack of expression for the indicated marker.

In agreement with the transcriptomic analysis, wholemount immunostaining of *Csf1r-iCre*;*Rosa*^*tdTom*^ mouse dermis at E15.5 identified TOM+ cells in dermal vessels with a lymphatic morphology that also expressed NRP2 and PROX1 (**Fig. 1E**). Such *Csf1r* lineage dermal LECs were already observed in *Csf1r-iCre*;*Rosa*^*tdTom*^ embryos at E13.5 (**Fig. 1F**). Quantification showed that TOM+ cells comprised 61.5 ± 9.0% and 64.0 ± 9.8% of PROX1+ cells in dermal lymphatic vasculature at E13.5 and E15.5, respectively (**Fig. 1F**). Notably, TOM+ cells were also observed in lymphatic vessels of the adult mouse ear dermis (**Fig. 1G**). Together, these data show that a subset of dermal LECs arises from a *Csf1r* cell lineage in the embryo and persists into adulthood.

### *Csf1r* lineage LECs do not arise from differentiated myeloid cells and lack endogenous *Csf1r* expression

*Csf1r* is a marker of differentiated myeloid cells (Byrne, Guilbert and Stanley, 1981; MacDonald *et al*., 2005; Nandi *et al*., 2012). To investigate whether TOM+ LECs arise from *Csf1r*-expressing myeloid cells, we introduced the *Csf1r-iCre*;*Rosa*^*tdTom*^ lineage trace into *Spi1* knockout mice, which lack the PU.1 transcription factor that is essential for the differentiation of haematopoietic progenitors into mature myeloid cells (Scott *et al*., 1997). Immunostaining for PROX1 and TOM showed that the proportion of *Csf1r* lineage LECs was similar in the dermis of E15.5 *Spi1*^*-/-*^ embryos and their wild-type littermates (**Fig. 2A**). The retention of *Csf1r* lineage LECs in *Spi1-*null mice demonstrates that these cells do not arise from differentiated myeloid cells and is reminiscent of the finding that PU.1 deficiency does not prevent the formation of *Csf1r* lineage BECs (Plein *et al*., 2018). In agreement with this idea, immunostaining of E17.5 dermis from *Spi1*^*-/-*^ mice carrying *Csf1r-iCre* and the alternative recombination reporter *Rosa*^*Yfp*^ for YFP identified *Csf1r* lineage ECs in both EMCN+ capillaries and the larger, NRP2+ lymphatic vessels (**Fig. 2B**). This finding also corroborates that lineage tracing of dermal LECs in the above studies was not specific to the *Rosa*^*tdTom*^ allele.

**Fig 2.**
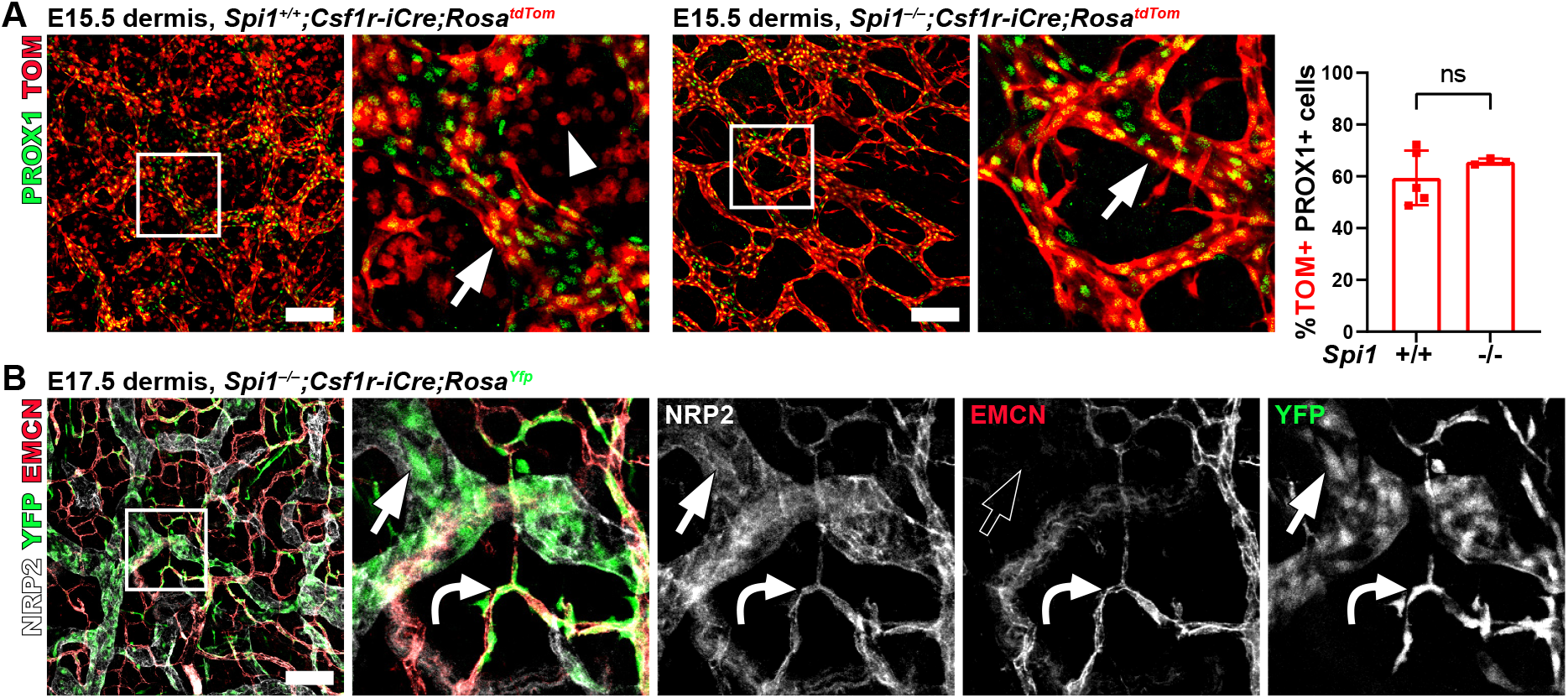
*Csf1r* lineage LECs persist in the absence of myeloid cells. **A** Representative immunofluorescence staining with the indicated markers and quantification of TOM+ LECs of E15.5 *Csf1r-iCre*;*Rosa*^*tdTom*^ dermis (scale bar: 100 μm) from mice with (*Spi1*^*+/+*^) or without (*Spi1*^*-/-*^) differentiated myeloid cells. The square indicates an area shown at higher magnification in the adjacent panel. The bar plot shows the fraction of TOM+ PROX1+ cells; mean ± SD, n = 5 *Spi1*^*-/-*^, n = 3 *Spi1*^*+/+*^ embryos; each dot represents the value from one embryo. **B** Immunofluorescence staining with the indicated markers of E17.5 *Spi1*^*-/-*^;*Csf1r-iCre*;*Rosa*^*Yfp*^ dermis (scale bar: 100 μm); the square indicates an area shown at higher magnification in the adjacent panel, and shown for the different markers also in grey scale. Arrows indicate TOM+ LECs; arrowheads indicate TOM+ macrophages; curved arrows indicate TOM+ BECs; empty symbols indicate lack of expression for the indicated marker.

Although our transcriptomic analysis suggested that E12.5 TOM+ ECs do not express *Csf1r* (**Fig. 1A,B**), this did not exclude that LECs arising at a later developmental stage to form organ lymphatic vasculature may begin to express *Csf1r*. To investigate whether *Csf1r* is actively expressed in dermal LECs, we used a *Csf1r-Egfp* transgenic expression reporter that faithfully reports endogenous *Csf1r* expression, including in macrophages (Sasmono *et al*., 2003). As expected, immunostaining of E15.5 *Csf1r-iCre*;*Rosa*^*tdTom*^;*Csf1r-Egfp* dermis identified EGFP expression in individual TOM+ cells with the morphology of tissue macrophages, but EGFP was not detected in TOM+ or TOM-LECs, which were identified as either NRP2+, PROX1+ or FLT4+ cells (**Fig. 3A**).

**Fig 3.**
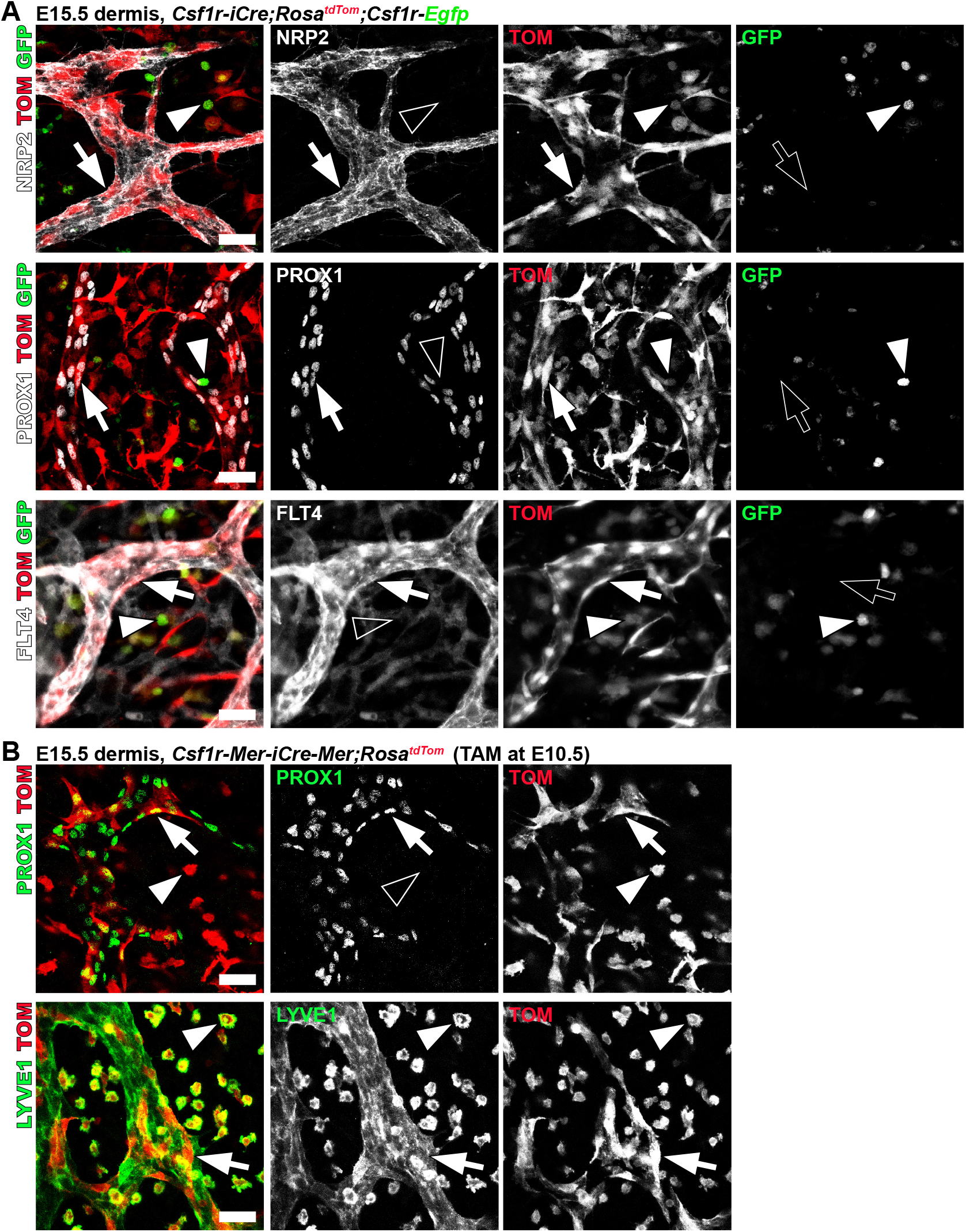
*Csf1r* lineage LECs do not themselves express *Csf1r* but derive from a *Csf1r*-expressing precursor. **A**,**B** Immunofluorescence staining with the indicated markers of *Csf1r-iCre*;*Rosa*^*tdTom*^;*Csf1r-Egfp* (**A**) and *Csf1r-Mer-iCre-Mer*;*Rosa*^*tdTom*^ mice injected with tamoxifen at E10.5 (**B**) (scale bars: 25 μm). Arrows indicate TOM+ LECs; arrowheads indicate TOM+ GFP+ (**A**) and TOM+ (**B**) macrophages; empty symbols indicate lack of expression for the indicated marker.

We further sought to corroborate that lineage tracing of dermal LECs in the above studies was not specific to the *Csf1r-iCre* transgene and could also be observed with an another *Csf1r*-mediated lineage trace. Thus, we combined the independently generated *Csf1r-Mer-iCre-Mer* transgene with the *Rosa*^*tdTom*^ reporter and administered tamoxifen at E10.5 to induce CRE activity, as previously done to corroborate the existence of *Csf1r* lineage BECs (Plein *et al*., 2018). Immunostaining for TOM and PROX1 or LYVE1 indeed identified TOM+ LECs in the dermis of E15.5 *Csf1r-Mer-iCre-Mer;Rosa*^*tdTom*^ embryos (**Fig. 3B**). Together, these results imply that a LEC subset arises from a PU.1-independent, *Csf1r*-expressing precursor population that is present in the E10.5 embryo.

### *Csf1r-iCre*-mediated *Prox1* targeting disrupts lymphatic vascular development

To investigate whether *Csf1r* lineage LECs are important for lymphatic development, we used conditional null floxed (fl) *Prox1* alleles (Iwano *et al*., 2012), in which CRE-mediated recombination of a stop codon preceding the *Egfp* gene under the control of the *Prox1* promoter results in EGFP expression and PROX1 inactivation (**Fig. 4A**). Wholemount immunostaining of E15.5 dermis from *Csf1r-iCre*;*Prox1*^*fl/+*^ mice with one copy of the mutant *Prox1* allele confirmed EGFP expression in LYVE1+ lymphatic vessels (**Fig. 4B**). Further, the *Csf1r-iCre-*mediated deletion of both functional *Prox1* alleles caused lymphatic phenotypes similar to, but milder than, those reported for global *Prox1* mutants (Wigle and Oliver, 1999; Johnson *et al*., 2008), including oedematous skin and blood-filled lymphatics (**Fig. 4C**). Specifically, 69% of E15.5 *Csf1r-iCre*;*Prox1*^*fl/fl*^ mutants presented with blood pools in the dermis, and 32% also had oedema under the dorsal skin and appeared “swollen” (**Fig. 4C,D**). Furthermore, immunostaining for the erythrocyte marker TER119 showed that control *Prox1*^*fl/fl*^ littermates with no *Cre* contained erythrocytes only within blood vessels, as expected (**Fig. 4E,F**), but that erythrocytes were present in both blood and lymphatic vessels of *Csf1r-iCre*;*Prox1*^*fl/fl*^ embryos lacking functional PROX1 (**Fig. 4E,F**). These phenotypes were also observed in hemizygous *Csf1r-iCre*;*Prox1*^*fl/+*^ embryos but with lower penetrance and severity (**Fig. 4F**). These experiments demonstrate that *Csf1r* lineage LECs make an essential contribution to lymphatic development.

**Fig 4.**
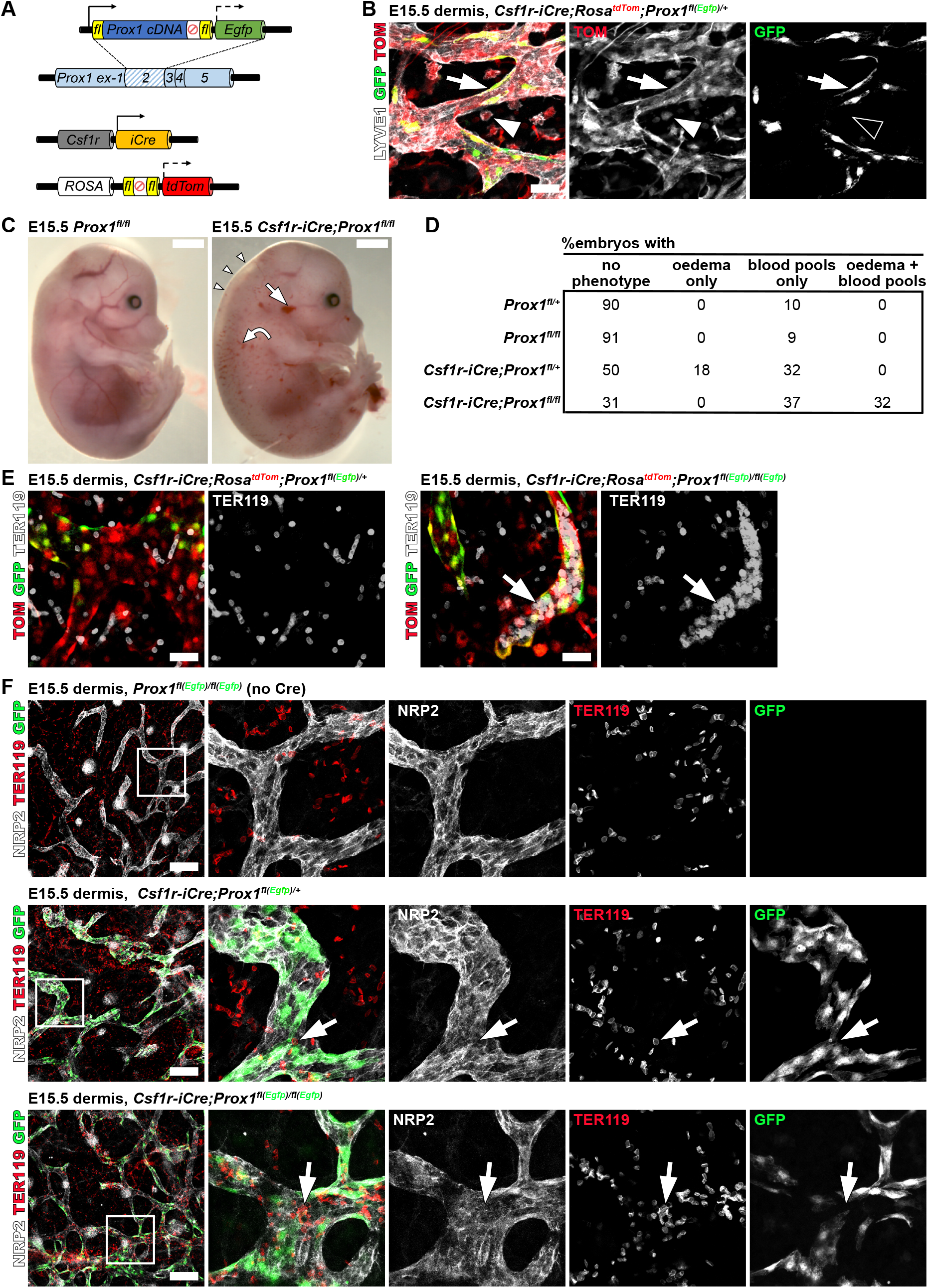
*Csf1r-iCre-mediated Prox1 targeting disrupts lymphatic vascular development*. **A**,**B** Strategy for the combined *Csf1r-iCre*-mediated lineage tracing and targeting of *Prox1* (**A**) and representative immunofluorescence staining with the indicated markers of E15.5 dermis from a heterozygously targeted *Csf1r-iCre*;*Prox1*^*fl(Egfp)/+*^ mouse (**B**) (scale bars: 25 μm). **C**,**D** Representative images of E15.5 *Prox1*^*fl/fl*^ embryos with or without *Csf1r-iCre* (scale bars: 1 mm) (**C**) and table showing the frequency of embryos displaying the indicated phenotype (**D**); n = 20 *Prox1*^*fl/+*^, n = 22 *Prox1*^*fl/fl*^, n = 22 *Csf1r-iCre*;*Prox1*^*fl/+*^, n = 19 *Csf1r-iCre*;*Prox1*^*fl/fl*^ from 11 litters. **E** Immunofluorescence staining with the indicated markers of E15.5 dermis of *Csf1r-iCre*;*Rosa*^*tdTom*^;*Prox1*^*fl(Egfp)/+*^ and *Csf1r-iCre*;*Rosa*^*tdTom*^;*Prox1*^*fl(Egfp)/fl(Egfp)*^ littermate mice identifies erythrocytes in mutant dermal lymphatic vessels that are co-labelled for TOM and GFP (scale bars: 100 μm). **F** Representative immunofluorescence staining with the indicated markers of E15.5 *Prox1*^*fl(Egfp)/fl(Egfp)*^ (no Cre, normal PROX1 function), *Csf1r-iCre*;*Prox1*^*fl(Egfp)/+*^ (heterozygous PROX1 deficiency) and E15.5 *Csf1r-iCre*;*Prox1*^*fl(Egfp)/fl(Egfp)*^ (homozygous PROX1 deficiency) dermis illustrates that TER119+ erythrocytes are located in NRP2+ lymphatic vessels of PROX1-deficient embryos. The square indicates an area shown at higher magnification in the adjacent panels and shown for the different markers also in grey scale (Scale bars: 100 μm). Arrows indicate TER119+ erythrocytes in lymphatic vessels.

## DISCUSSION

According to the classic model of lymphatic development, LECs arise from *Tie2* lineage venous BECs following induction of *Prox1* expression (Wigle and Oliver, 1999; Srinivasan *et al*., 2007; Yang *et al*., 2012; Jafree *et al*., 2021). Since, other lineage tracing studies have revealed organ-specific LEC contributions from non-venous sources, including paraxial mesoderm, second heart field derivatives and haemogenic endothelium in the heart, dermis and/or mesentery (Srinivasan *et al*., 2007; Klotz *et al*., 2015; Stanczuk *et al*., 2015; Ulvmar and Mäkinen, 2016; Maruyama *et al*., 2019; Stone and Stainier, 2019; Lioux *et al*., 2020; Klaourakis, Vieira and Riley, 2021). Our results add to these findings by demonstrating that a significant proportion of dermal LECs arises from a *Csf1r-*expressing cell lineage that is important for lymphatic development. Firstly, transcriptomic analysis revealed enrichment of canonical lymphatic markers in *Csf1r* lineage traced ECs at E12.5, including *Prox1, Flt4* and *Nrp2*. Secondly, wholemount immunostaining demonstrated the presence of *Csf1r* lineage LECs in the embryonic dermis already from early stages of lymphatic network formation in this tissue. Thirdly, *Csf1r* lineage LECs make a non-redundant contribution to lymphatic development, demonstrated by lymphatic defects in mice with the *Csf1r-iCre*-mediated deletion of *Prox1*, including dermal oedema and blood-filled dermal lymphatic vessels.

The phenotypes caused by the *Csf1r-iCre*-mediated deletion of *Prox1* were similar to, but milder than, the lymphatic defects reported for global or pan-endothelial *Prox1* mutants (Wigle and Oliver, 1999; Johnson *et al*., 2008). The milder severity of *Csf1r* lineage *Prox1* mutants compared to global or pan-endothelial *Prox1* mutants is consistent with our observation that not all LECs can be assigned to the *Csf1r* lineage based on observations with a *Csf1r-iCre*-induced recombination reporter. Thus, only 60% of dermal LECs are targeted by *Csf1r-iCre*, consistent with the existence of other LEC progenitor sources previously reported for the dermis (Martinez-Corral *et al*., 2015; Stone and Stainier, 2019).

We considered several possibilities to explain how LECs may arise from a *Csf1r* lineage. On the one hand, it is conceivable that *Csf1r* is expressed in LEC themselves. However, two independent pieces of evidence argue against this possibility. First, E12.5 ECs did not contain detectable levels of *Csf1r* transcripts. Second, E15.5 dermal LECs did not appear to express the *Csf1r-Egfp* reporter that was previously shown to recapitulate endogenous *Csf1r* expression. On the other hand, *Csf1r* is a key gene expressed in myeloid cells and macrophages, raising the possibility that these cells might transdifferentiate into LECs that contribute to dermal lymphatics. However, *Csf1r* lineage LECs were retained in *Spi1*-deficient embryos, which lack differentiated myeloid cells (Scott *et al*., 1997). These findings are analogous to our previous observations for the blood vasculature, whereby *Csf1r* lineage BECs did not express *Csf1r* and persisted independently of myeloid differentiation (Plein *et al*., 2018). Therefore, our results are consistent with a model in which *Csf1r* lineage LECs arise from a PU.1-independent progenitor that transiently expresses *Csf1r* before adopting a lymphatic endothelial fate, either directly via lymphvasculogenesis or by differentiating first into BECs that then specialise into LECs in the dermis. Both concepts are compatible with previous reports suggesting that dermal lymphatics derive from LEC clusters (Martinez-Corral *et al*., 2015) and blood vascular capillaries (Pichol-Thievend *et al*., 2018)

The underlying reasons for cooperation of different LEC sources to build a functional lymphatic network in the dermis remain unknown, as is knowledge of whether lineage heterogeneity contributes to regional specialisation or adaptive responses during lymphatic growth and repair. In the future, the discovery of alternative lymphatic progenitor sources may inform strategies to stimulate lymphangiogenesis in pathological contexts such as lymphoedema, chronic inflammation or cardiac disease. In particular, further work to identify the signals that govern the specification and expansion of different LEC lineages from distinct progenitors might open new therapeutic avenues, for example, via directed differentiation of induced pluripotent stem cells for personalised medicine into cells that can give rise to organ-specific lymphatic vasculature.

## ACKNOWLEDGEMENTS

We thank the staff of the Biological Resources, FACS and Imaging Facilities at the UCL Institute of Ophthalmology for their technical support. This research was supported by grants from the British Heart Foundation to RC and CR [FS/19/14/34170] and CR [PG23/11301] and an investigator award from the Wellcome to CR [205099/Z/16/Z].

## AUTHORSHIP CONTRIBUTIONS

GC, RC, AP, AF and CR conceived and designed the study. GC and CR co-wrote the manuscript. GC, RC, AP and LD performed experiments. AF performed bioinformatic analyses. CR supervised the project. All authors read and approved the submitted manuscript.

## DISCLOSURE OF CONFLICTS OF INTEREST

The authors declare that they have no competing interests.

## Notes

### Competing Interest Statement

The authors have declared no competing interest.

### Summary of Updates

The figures were accidentally duplicated in the previous version. This has now been corrected.

## REFERENCES

Alitalo, K. (2011) ‘The lymphatic vasculature in disease’, Nature medicine, 17(11), pp. 1371–1380.

Banerji, S. et al. (1999) ‘LYVE-1, a new homologue of the CD44 glycoprotein, is a lymph-specific receptor for hyaluronan’, The Journal of cell biology, 144(4), pp. 789–801.

Bolger, A.M., Lohse, M. and Usadel, B. (2014) ‘Trimmomatic: a flexible trimmer for Illumina sequence data’, Bioinformatics (Oxford, England), 30(15), pp. 2114–2120.

Burnett, S.H. et al. (2004) ‘Conditional macrophage ablation in transgenic mice expressing a Fas-based suicide gene’, Journal of leukocyte biology, 75(4), pp. 612–623.

Byrne, P. V., Guilbert, L.J. and Stanley, E.R. (1981) ‘Distribution of cells bearing receptors for a colony-stimulating factor (CSF-1) in murine tissues’, The Journal of cell biology, 91(3 Pt 1), pp. 848–853.

Deng, L. et al. (2010) ‘A novel mouse model of inflammatory bowel disease links mammalian target of rapamycin-dependent hyperproliferation of colonic epithelium to inflammation-associated tumorigenesis’, American Journal of Pathology, 176(2), pp. 952–967.

Dobin, A. et al. (2013) ‘STAR: ultrafast universal RNA-seq aligner’, Bioinformatics (Oxford, England), 29(1), pp. 15–21.

François, M. et al. (2012) ‘Segmental territories along the cardinal veins generate lymph sacs via a ballooning mechanism during embryonic lymphangiogenesis in mice’, Developmental Biology, 364(2), pp. 89–98.

Iwano, T. et al. (2012) ‘Prox1 postmitotically defines dentate gyrus cells by specifying granule cell identity over CA3 pyramidal cell fate in the hippocampus’, Development, 139(16), pp. 3051–3062.

Jafree, D.J. et al. (2021) ‘Mechanisms and cell lineages in lymphatic vascular development’, Angiogenesis 2021 24:2, 24(2), pp. 271–288.

Johnson, N.C. et al. (2008) ‘Lymphatic endothelial cell identity is reversible and its maintenance requires Prox1 activity’, Genes & Development, 22(23), pp. 3282–3291.

Klaourakis, K., Vieira, J.M. and Riley, P.R. (2021) ‘The evolving cardiac lymphatic vasculature in development, repair and regeneration’, Nature Reviews Cardiology 2021 18:5, 18(5), pp. 368–379.

Klotz, L. et al. (2015) ‘Cardiac lymphatics are heterogeneous in origin and respond to injury’, Nature 2015 522:7554, 522(7554), pp. 62–67.

Liao, Y., Smyth, G.K. and Shi, W. (2014) ‘featureCounts: an efficient general purpose program for assigning sequence reads to genomic features’, Bioinformatics (Oxford, England), 30(7), pp. 923–930.

Lioux, G. et al. (2020) ‘A Second Heart Field-Derived Vasculogenic Niche Contributes to Cardiac Lymphatics’, Developmental Cell, 52(3), pp. 350–363.e6.

MacDonald, K.P.A. et al. (2005) ‘The colony-stimulating factor 1 receptor is expressed on dendritic cells during differentiation and regulates their expansion’, Journal of immunology (Baltimore, Md. : 1950), 175(3), pp. 1399–1405.

Madisen, L. et al. (2010) ‘A robust and high-throughput Cre reporting and characterization system for the whole mouse brain’, Nature neuroscience, 13(1), pp. 133–140.

Martin-Almedina, S., Mortimer, P.S. and Ostergaard, P. (2021) ‘Development and physiological functions of the lymphatic system: insights from human genetic studies of primary lymphedema’, Physiological reviews, 101(4), pp. 1809–1871.

Martinez-Corral, I. et al. (2015) ‘Nonvenous origin of dermal lymphatic vasculature’, Circulation Research, 116(10), pp. 1649–1654.

Maruyama, K. et al. (2019) ‘Isl1-expressing non-venous cell lineage contributes to cardiac lymphatic vessel development’, Developmental Biology, 452(2), pp. 134–143.

Mayeuf-Louchart, A. et al. (2014) ‘Notch regulation of myogenic versus endothelial fates of cells that migrate from the somite to the limb’, Proceedings of the National Academy of Sciences of the United States of America, 111(24), pp. 8844–8849.

McKercher, S.R. et al. (1996) ‘Targeted disruption of the PU.1 gene results in multiple hematopoietic abnormalities’, The EMBO Journal, 15(20), p. 5647.

Nandi, S. et al. (2012) ‘The CSF-1 receptor ligands IL-34 and CSF-1 exhibit distinct developmental brain expression patterns and regulate neural progenitor cell maintenance and maturation’, Developmental Biology, 367(2), pp. 100–113.

Oliver, G. (2022) ‘Lymphatic endothelial cell fate specification in the mammalian embryo: An historical perspective’, Developmental Biology, 482, pp. 44–54.

Pichol-Thievend, C. et al. (2018) ‘A blood capillary plexus-derived population of progenitor cells contributes to genesis of the dermal lymphatic vasculature during embryonic development’, Development (Cambridge, England), 145(10).

Plein, A. et al. (2018) ‘Erythro-myeloid progenitors contribute endothelial cells to blood vessels’, Nature 2018 562:7726, 562(7726), pp. 223–228.

Qian, B.Z. et al. (2011) ‘CCL2 recruits inflammatory monocytes to facilitate breast-tumour metastasis’, Nature 2011 475:7355, 475(7355), pp. 222–225.

Sasmono, R.T. et al. (2003) ‘A macrophage colony-stimulating factor receptor–green fluorescent protein transgene is expressed throughout the mononuclear phagocyte system of the mouse’, Blood, 101(3), pp. 1155–1163.

Scott, E.W. et al. (1997) ‘PU.1 furactions in a cell-autonomous manner to control the differentiation of multipotential lymphoid-myeloid progenitors’, Immunity, 6(4), pp. 437– 447.

Srinivas, S. et al. (2001) ‘Cre reporter strains produced by targeted insertion of EYFP and ECFP into the ROSA26 locus’, BMC Developmental Biology 2001 1:1, 1(1), pp. 4–.

Srinivasan, R.S. et al. (2007) ‘Lineage tracing demonstrates the venous origin of the mammalian lymphatic vasculature’, Genes & development, 21(19), pp. 2422–2432.

Stanczuk, L. et al. (2015) ‘CKit lineage hemogenic endothelium-derived cells contribute to mesenteric lymphatic vessels’, Cell Reports, 10(10), pp. 1708–1721.

Stone, O.A. and Stainier, D.Y.R. (2019) ‘Paraxial Mesoderm Is the Major Source of Lymphatic Endothelium’, Developmental Cell, 50(2), pp. 247–255.e3.

Ulvmar, M.H. and Mäkinen, T. (2016) ‘Heterogeneity in the lymphatic vascular system and its origin’, Cardiovascular Research, 111(4), pp. 310–321.

Varet, H. et al. (2016) ‘SARTools: A DESeq2- and EdgeR-Based R Pipeline for Comprehensive Differential Analysis of RNA-Seq Data’, PloS one, 11(6).

Veikkola, T. et al. (2001) ‘Signalling via vascular endothelial growth factor receptor-3 is sufficient for lymphangiogenesis in transgenic mice’, The EMBO journal, 20(6), pp. 1223– 1231.

Wigle, J.T. et al. (2002) ‘An essential role for Prox1 in the induction of the lymphatic endothelial cell phenotype’, The EMBO journal, 21(7), pp. 1505–1513.

Wigle, J.T. and Oliver, G. (1999) ‘Prox1 function is required for the development of the murine lymphatic system’, Cell, 98(6), pp. 769–778.

Yang, Y. et al. (2012) ‘Lymphatic endothelial progenitors bud from the cardinal vein and intersomitic vessels in mammalian embryos’, Blood, 120(11), pp. 2340–2348.

Yuan, L. et al. (2002) ‘Abnormal lymphatic vessel development in neuropilin 2 mutant mice’, Development (Cambridge, England), 129(20), pp. 4797–4806.

